# The MYC antagonist MNT autoregulates its expression and supports proliferation in MAX deficient cells

**DOI:** 10.1101/212944

**Authors:** M Carmen Lafita-Navarro, Judit Liaño-Pons, Andrea Quintanilla, Ignacio Varela, Fabiana Ourique, Gabriel Bretones, Rosa Blanco, Julia Aresti, Patrick Carroll, Peter Hurlin, Robert N. Eisenman, M. Dolores Delgado, Javier León

## Abstract

MNT is a transcription factor of the MXD family. MNT-MAX dimers down-regulate genes by binding to E-box sequences, which can also be bound by MYC-MAX to activate transcription. MNT has been described as a modulator of MYC activity but little is known about MNT regulation and whether MNT has MAX-independent functions. Using a MAX deficient cell line and siRNA-mediated silencing of MAX, we show that in the absence of MAX, the total MNT levels are elevated and that MNT localizes both in the cytoplasm and the nucleus. In contrast, MNT is predominantly nuclear when MAX is expressed. MNT is required for optimal cell proliferation even in the absence of MAX, being the first report of a MAX-independent function of MNT. Interestingly, MNT forms homodimers and autoregulates its expression by repressing its own promoter. The tight MNT regulation and its activity in absence of MAX suggest its importance on cell homeostasis.

## INTRODUCTION

MNT (also called MXD6) is a basic helix-loop-helix leucine zipper (bHLHLZ) protein and it is the most divergent member of the MXD family, that also includes MXD1, MXI1, MXD3 and, MXD4. MNT forms heterodimers with MAX through the bHLHLZ domain, and binds to E‐box DNA sequences. MYC is one of the most prevalent human oncoproteins. MYC can also interact with MAX and bind E-boxes (*Conacci-Sorrell et al 2014, Dang 2012*). Whereas MYC-MAX, upon binding to E-boxes, acts primarily as transcriptional activator, the typical effect of MNT-MAX is the transcriptional repression (*Hurlin et al 1997, Meroni et al 1997*). MNT can bind not only to MAX but also to the MAX-like HLH protein MLX (*Billin et al 1999, Meroni et al 2000*) which alternatively can interact with MLXIP (MONDOA) and MLXIPL (MONDOB) proteins. Therefore, MNT participates both in the MAX- and MLX-centred networks [reviewed in (*Diolaiti et al 2015, Yang & Hurlin 2017*)] serving as the link between the MAX-MYC and MLX-MONDO networks.

MNT is expressed constitutively in proliferating and quiescent cells and protein levels do not show major fluctuations when quiescent cells are mitotically stimulated (*Hurlin et al 1997, Hurlin et al 2003, Walker et al 2005*). MNT^−/−^ mice are not viable (*Hurlin et al 2003, Toyo-oka et al 2004*) while MXD1^-/-^, MXI1^-/-^ and MXD3^-/-^ mice survive, suggesting that MNT function is not redundant with that of the other MXD proteins (*Foley et al 1998, Queva et al 1999, Schreiber-Agus & DePinho 1998*). Moreover, MNT is the only MXD protein in invertebrates (*Diolaiti et al 2015*).

Consistent with MNT functioning as a MYC transcriptional antagonist, enforced MNT expression inhibits cell proliferation and impairs MYC-dependent transformation (*Hurlin et al 1997, Walker et al 2005*). The deficiency or down‐regulation of MNT in fibroblasts leads to increased proliferation (i.e., similarly to MYC-overexpression) and partially rescues the proliferative arrest caused by MYC deficiency (*Dezfouli et al 2006, Hurlin et al 2003, Nilsson et al 2004, Walker et al 2005*). MNT ablation *in vivo* leads to breast and T-cell tumors (*Dezfouli et al 2006, Hurlin et al 2003, Toyo-oka et al 2004*), and partial or total MNT deficiency in mice models impairs MYC-dependent tumorigenesis (*Link et al 2012; Campbell et al 2017),* suggesting that *MNT* may function as a tumor suppressor gene. On the other hand, *MNT* knock-out in some cell models inhibits proliferation and promotes apoptosis (*Dezfouli et al 2006, Hurlin et al 2003, Nilsson et al 2004*). Thus, as MYC and MNT coexist in proliferating cells, it has been suggested a role of MNT as a “buffer” of MYC activity.

In the present work we studied possible MAX-independent functions of MNT using as the main model UR61 cells. These rat pheochromocytoma cells that do not express a functional MAX protein but a truncated form (termed MAX^PC12^) that lacks the second helix and leucine zipper region of the bHLH-LZ domain which are the regions responsible for dimerization with MYC and MNT (*Hopewell & Ziff 1995*). Here we describe a change of MNT subcellular localization depending on MAX expression and that MNT is required for optimal cell proliferation even in the absence of MAX, being the first example of a MAX-independent function of MNT. Finally, we show that MNT transrepresses its own promoter and MNT ability to regulate gene expression in the absence of MAX.

## RESULTS

### MNT levels depend on MAX

We first studied MNT levels in proliferating cells from eleven cell lines derived from different tissues and species, including two cell lines lacking MAX: UR61 and human small cell lung carcinoma H1417 cells (*Romero et al 2014*). The results showed that, although MNT expression varies among cell lines, the two MAX-deficient cell lines expressed high MNT levels (Figure 1A). MNT was expressed in all cell lines as a protein doublet, due to a slower migrating phosphorylated MNT form (*Popov et al 2005*).

**Figure 1.**
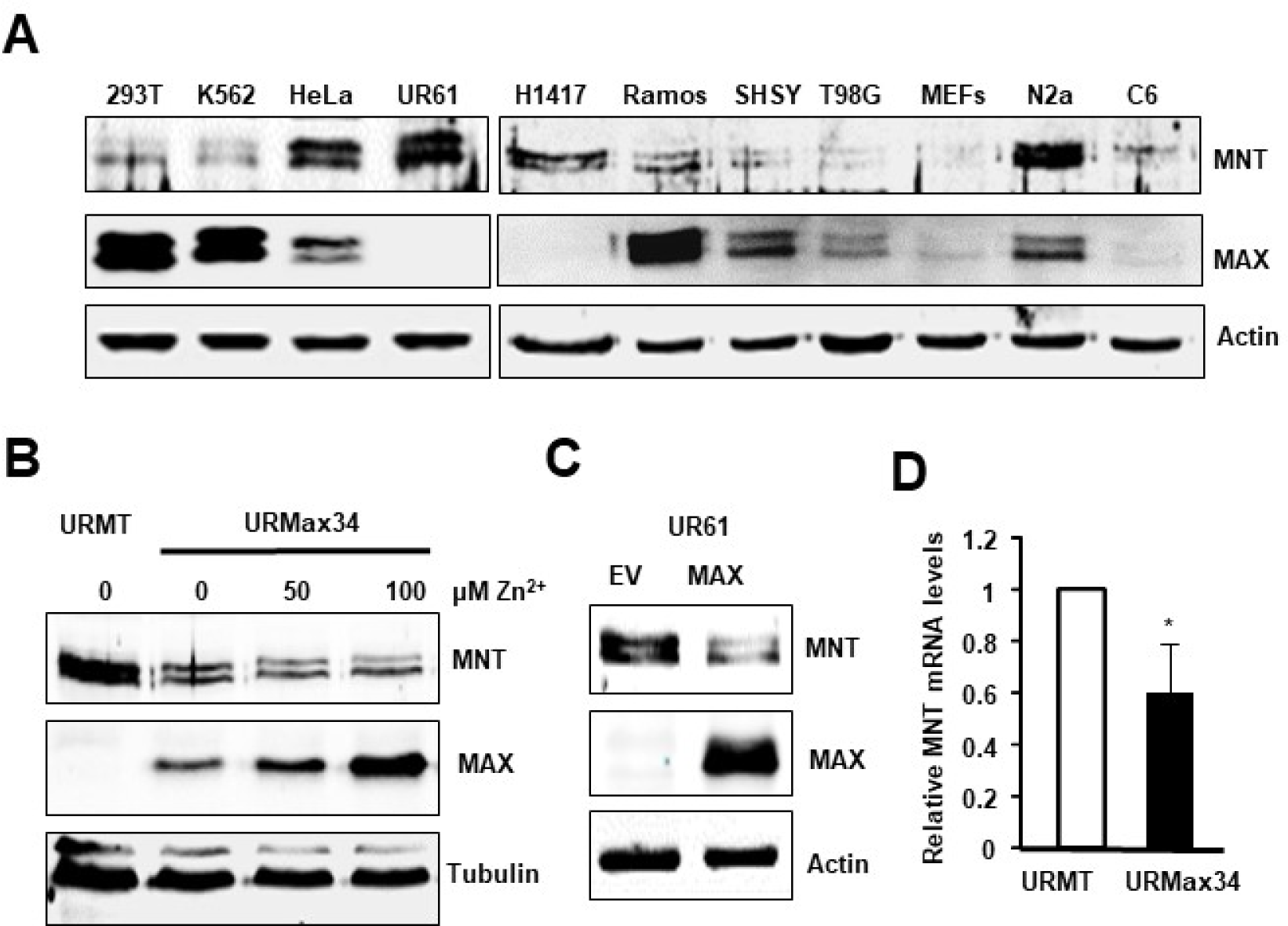
High MNT ex pression in MAX-deficient cells. (A) Cell Iysates of the indicated cell lines were analyzed by immunoblot to determine the levels of MNT and MAX. β-actin levels were determined for protein loading control. The MAX-deficient cell Iines anaIyzed were: UR61 (rat pheochromocytoma) and H1417 (human small cell lung carcinoma). The rest are MAX-expressing cells: HEK293T (human embry onic kidney, 293T), K 562 (human chronic myeloid leukaemia), HeLa (human cervical cancer), Ramos (human B cell lymphoma), SH-SY-Y5 (SHSY, human neuroblastoma), T98G (human glioblastoma), MEFs (mouse embry onic fibroblasts), Neuro-2a (N2a, mouse neuroblastoma) and C6 (rat glioma). (B) Control URMT and URMax34 cells were treated for 24 h with 50 and 100 μM Zn^+2^ and the MNT and MAX protein expression were determined by immunoblot. α-tubulin levels were determined as a control for protein loading. The experiment was repeated three times. (C) Levels of MNT and MAX determined by immunoblot in UR61 cells 24 h after transfection with a MAX expression vector or the empty vector pCEFL (EV). β-actin levels are shown as a protein loading control. The result was reproduced in three experiments. (D) mRNA expression determined by RT-qPCR in URMT and URMax34 cells treated for 24 h with 100 μM Zn^+2^. Data are represented as mean ±s.d. from three independent experiments, *\*P* = 0.012.

The high MNT expression in the two MAX-deficient cell lines led us to explore whether MAX influenced MNT levels. For this purpose, we transfected UR61 cells with a construct carrying human MAX cDNA driven by the metallothionein promoter, which is activated by Zn^2+^ cations (*Canelles et al 1997*). Several clones were isolated and two of them with robust MAX induction were mixed and the resulting cell line was termed URMax34. We also generated a cell line transfected with the empty vector, termed URMT. The induction of MAX in response to Zn^2+^ in URMax34 cells was confirmed by immunoblot (Figure 1B). We examined the effect of MAX induction on MNT levels in URMax34 cells. The results showed that MNT was down-regulated upon MAX induction by Zn^2+^ (Figure 1B). To confirm this result and rule out effects potentially related to the generation of stably transfected clones (as URMax34 system), UR61 cells were transiently transfected with a MAX expression vector and the results showed a decrease in MNT protein levels in MAX-transfected cells (Figure 1C). It is noteworthy that the r-expression of MAX in these cells provoked the down-regulation of MNT at the mRNA levels, as determined by RT-qPCR (Figure 1D).

We sought to confirm this in a different cell type and species. For this purpose, a MAX expression vector was transfected into human myeloid leukemia K562 cells. The immunoblot results showed lower MNT levels in the cells with MAX overexpression (Figure 2A). Next, we used the Kmax12 cell line (*Canelles et al 1997*), a K562 derivative carrying a MAX transgene which expression is induced by Zn^2+^. Induction of MAX expression in Kmax12 cells resulted in a concomitant MNT downregulation (Figure 2B). We also used the opposite approach, i.e. depleting cells of MAX and analyzing the expression of MNT. As shown in Figure 2C, MNT protein expression was upregulated when MAX was silenced with siRNA in K562. Interestingly, MNT mRNA was also up-regulated in MAX-silenced cells (Figure 2D). Altogether, the results showed that low MAX levels result in MNT up-regulation, likely at the transcriptional level.

**Figure 2.**
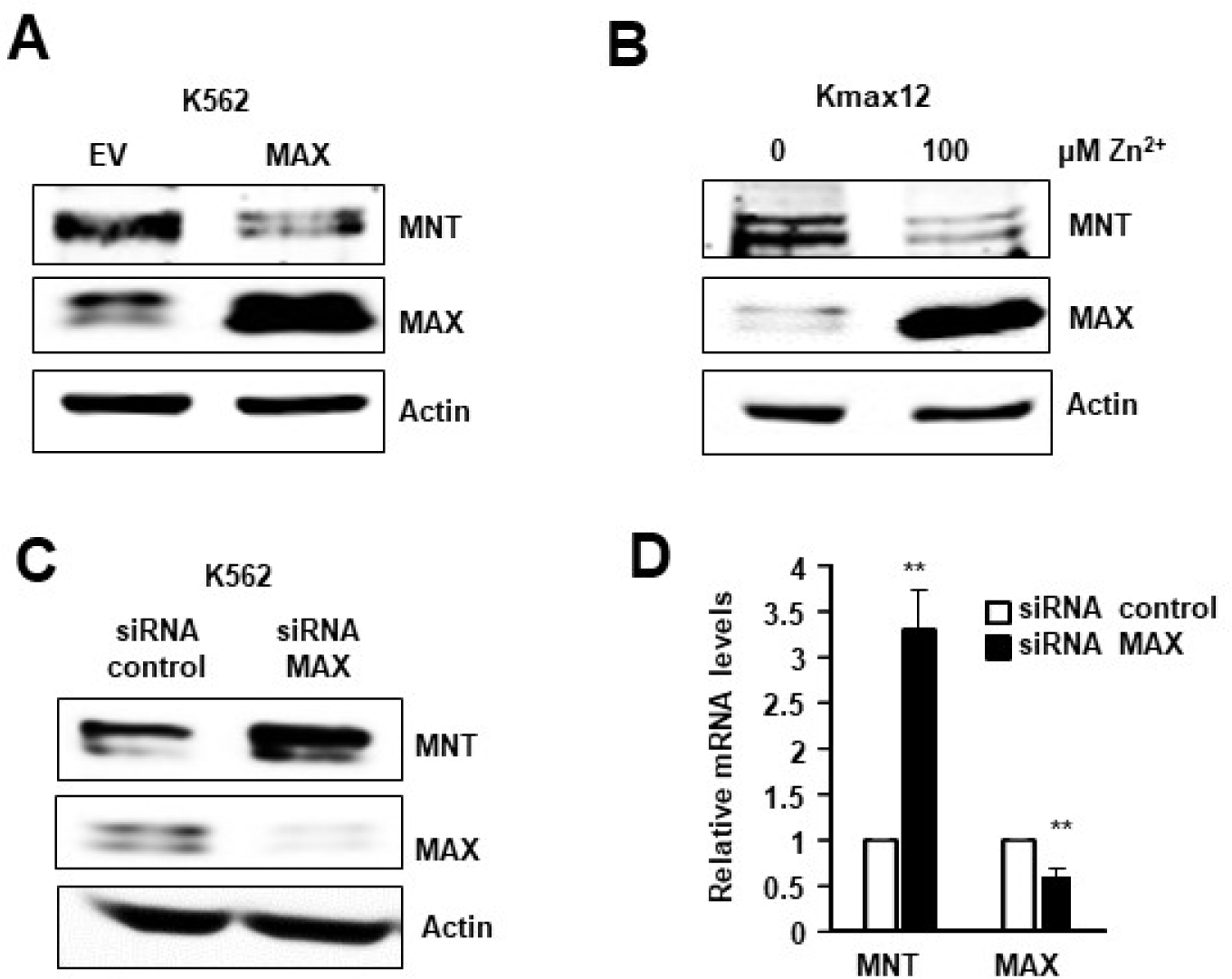
MAX regulates MNT expression in K562 cells. (A) MNT and MAX levels determined by imm unoblot in K562 cells 24 h after transfection with a MAX expression vector. The results were reproduced in three biological replicates (B) MNT and MAX levels determined by immunoblot in Kmax12 cells treated for 48 h with 100 lJM Zn^2+^ to induce MAX expression. The results were reproduced in three biological replicatess. (C) MNT and MAX protein expression analyzed by immunoblot in K562 48 h after transfection with siRNA against *MAX* gene. β-actin levels are shown as a protein loading control. The results were reproduced in two experi ments (D) MNT and MAX mRNA expression analyzed by RT-qPCR 48 h after transfection with siRNA against *MAX* gene. Data are shown as mean ± s.d. of three biological replicates. *\*P <* 0.005

### MNT binds and repress its own promoter in the presence of MAX

The above results showing that MNT down-regulation took place at the mRNA level prompted us to investigate if MNT impairs the activity of its own promoter. Bioinformatic analysis of human, mouse and rat MNT promoter regions revealed that there are two E-box sequences within 1 kb upstream the transcriptional start site of *MNT* (one canonical E-box, CACGTG (E-box 1), and one non-canonical, CATGTG (E-box 2) that are conserved among these three different species (Figure 3A).

**Figure 3.**
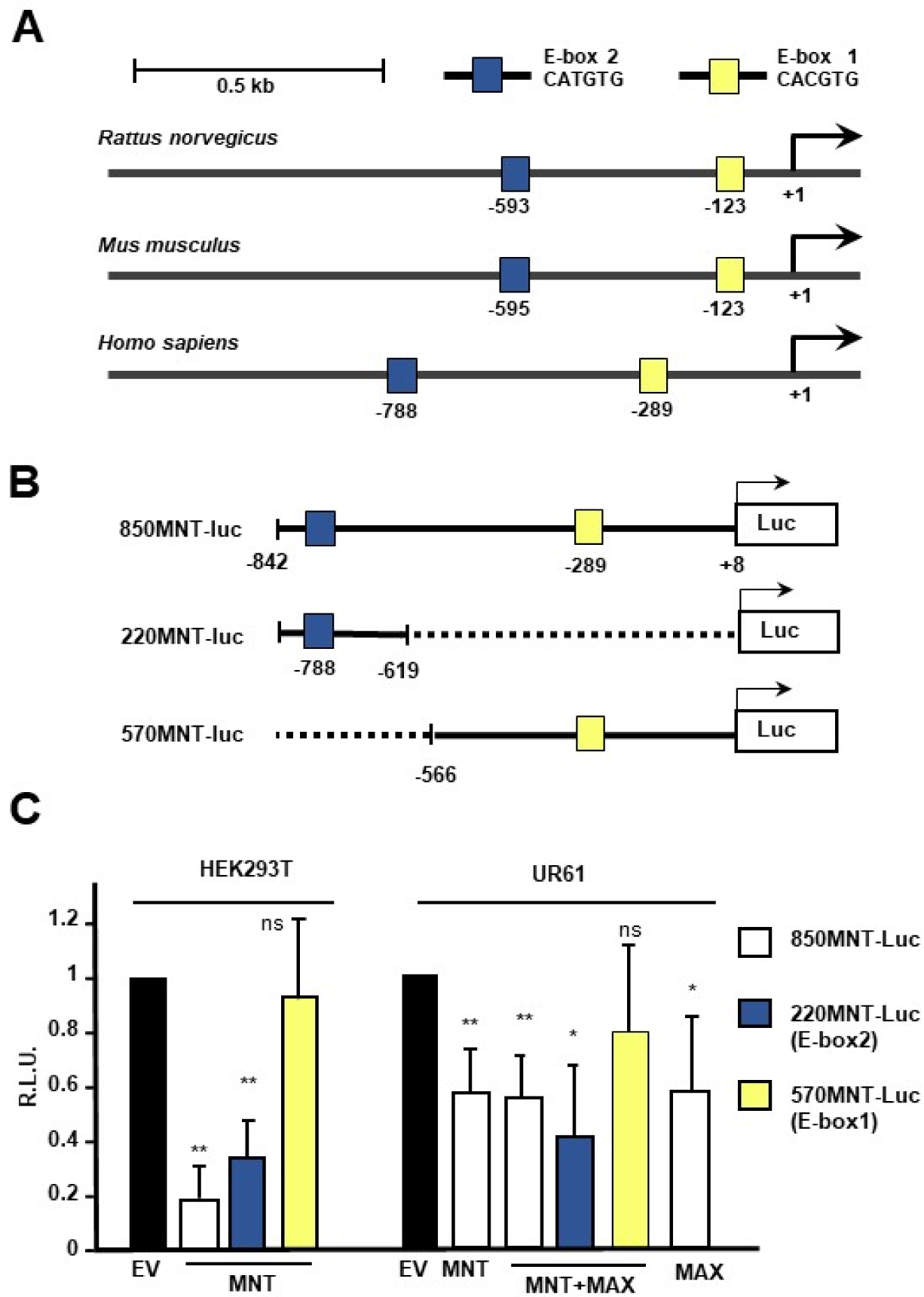
MNT regulates its own promoter. (A) Schematic representation of human, rat and mouse MNT promoters showing two conserved E-bo xes. The coordinates correspond to the 5′ nucleotide of each E-box, using the UCSC genome browser (http://genome.ucsc.edu/, release GRCh37/hg 19). (B) Schematic representation of luciferase reporters driven by human MNT promoter generated in this work. The longest reporter (850 bp upstream the transcription start site) contains two conserved E-boxes. The coordinates are as in USCS browser (release GRCh37/hg19). (C) Luciferase assays in the indicated cell lines transfected with the reporters described above plus MNT and/or MAX expressing vectors or the corresponding empty vectors (EV). Luciferase activity was assayed 36 h or 24 h after transfection of UR6 1 and HEK293T cells, respectively. The data are shown as mean ± s.e.m. of 9 (for 850MNT-luc) or 4 independent transfections (for the rest of experimental points). \*\**P* < 0.005; ~\**P* < 0.01.

We then constructed a luciferase reporter carrying the 850 bp upstream region of the transcription start site from the human MNT gene. The construct was termed 850MNT-Luc (Figure 3B). HEK293T cells (which express MAX) were transfected with the 850MNT-Luc and MNT expression vectors. The results showed that MNT overexpression led to a reduction in the luciferase activity (Figure 3C), suggesting that MNT-MAX negatively regulates the MNT promoter. To determine the contribution of the two E-boxes in the MNT-mediated negative autoregulation we constructed two reporters containing each of the E-boxes, termed 220MNT-Luc (containing the last 220 bp of the 850MNT-Luc reporter which includes the E-box 2) and 570MNT-Luc (containing the first 570 bp of the 850MNT-Luc reporter which includes the E-box 1) (Figure 3B). The luciferase assays in HEK293T cells showed that the E-box 2, mapping at -788 was critical for MNT-mediated downregulation of its own promoter (Figure 3C, left panel).

We also investigated the activity of the MNT promoter in UR61 cells, which do not express MAX. UR61 cells were transfected with the 850MNT-Luc vector together with MNT and MAX expression vectors. The results also showed a decrease in the luciferase activity although less than in HEK293T cells (Figure 3C, right panel). The expression of MNT alone also led to a decrease in the luciferase activity, suggesting that MNT can downregulate MNT promoter in UR61 cells in the absence of MAX. The repressive effect of MNT was stronger in HEK293T cells than in UR61 cells, which may be explained by the limited overexpression of MNT protein achieved in transfected UR61 cells (shown below). Co-transfection of MNT and MAX resulted in a decrease in the activity of both promoter constructs (850MNT-Luc and 220MNT-Luc, carrying fragments of the human MNT promoter of 842 and 223 bp, respectively). Since MNT-MAX dimers bind E-boxes in the promoters to repress transcription, we analysed the ChIP-seq data published by the ENCODE project (http://genome.ucsc.edu/ENCODE).

The data revealed two regions bound by MAX in the human *MNT* promoter that encompass the two E-boxes (Figure 4A). We also analysed the ChIP peaks for MAX, MYC and MXI1 proteins on the *MNT* promoter in other human cell lines as H1ES, HeLa, NB4, A549, GM78, HEPG, SKSM and IMR90. The analysis revealed that the three proteins presented peaks at the same positions near the transcription start site of the human MNT gene (Figure 4*-figure supplement 1*), suggesting that members of the MYC and MXD family bind and possibly regulate the promoter of *MNT*.

**Figure 4.**
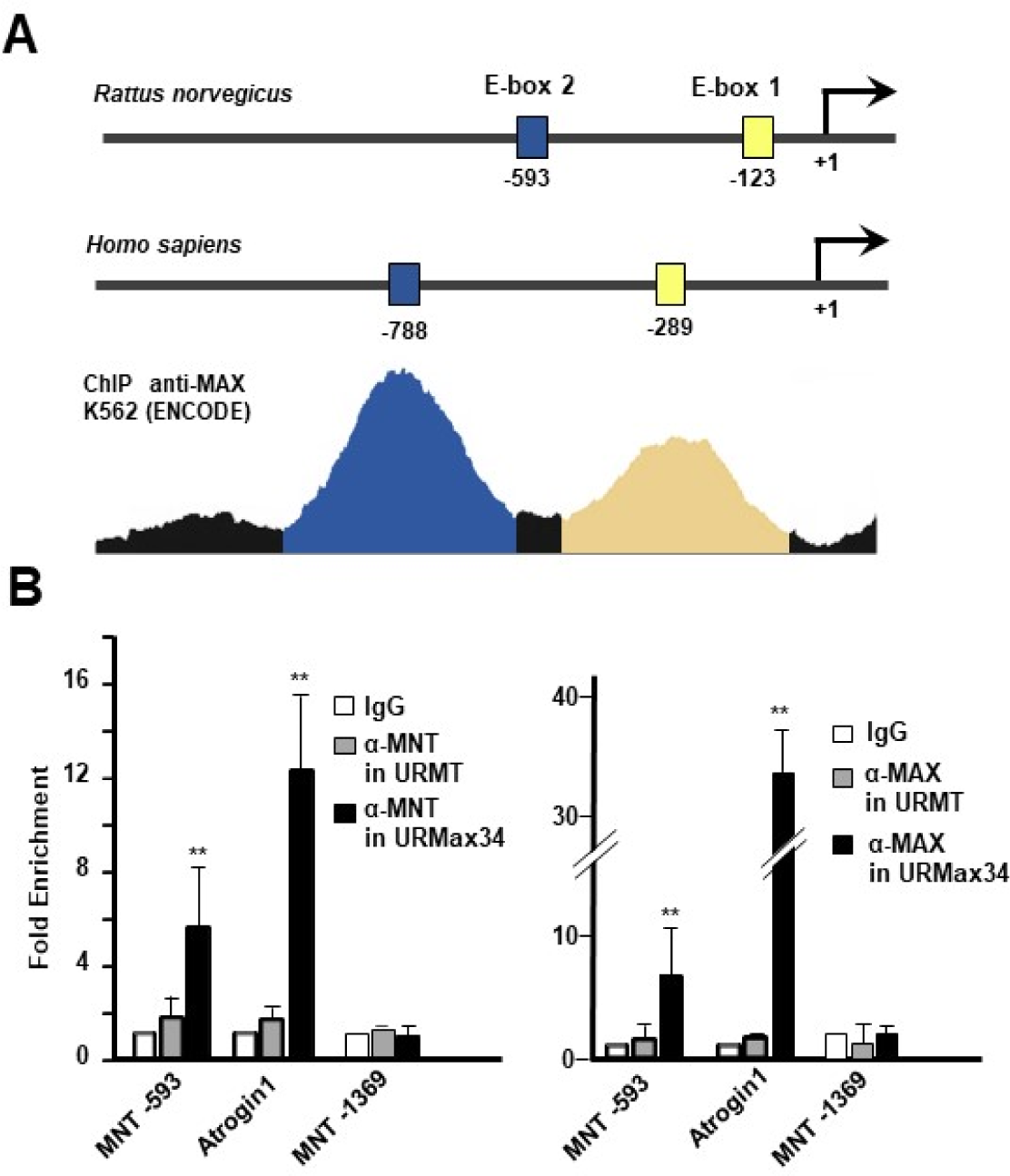
MNT binds to *MNT* promoter. (A) Schematic representation of human and rat MNT promoters showing the peaks for MAX of the ChIP experiments with anti-MAX antibody on human K562 cell line as published by the ENCODE project. (B) ChIP with anti-MNT (left) and anti-MAX (right) antibodies on the MNT promoter of URMT and URMax34 cells, both treated with 100 μM Zn^2+^ for 24 h. MNT and MAX binding to *MNT* gene promoter in the region containing the E-box 2 (-59 3 in rat gene) was studied by qPCR Amplicons from Atrogin-1 (*FBX032*) were used as positive control for MNT binding and a region 1389 bp upstream from the TSS without E-boxes was used as negative control. The data are means ±s.e.m. from four independent experiments. \**P* < 0.05.

As MNT was downregulated when MAX was re-expressed, we hypothesized that MNT-MAX heterodimers might bind to the *MNT* promoter and down-regulate its own expression. To explore this hypothesis, a ChIP assay was performed with MNT and MAX antibodies in URMT and URMax34 treated with Zn^2+^ to induce MAX. We studied the -593 E-box of the *MNT* promoter, as well as a region of the *Atrogin-1 (FBXO32*) promoter as a positive control for MNT binding (*Terragni et al 2011*). A region mapping at 1369 bp upstream on MNT transcription start site with no E-boxes was used as negative control. As shown in Figure 4B, MNT and MAX bound to the *MNT* promoter in URMax34 cells. However, in the MAX-deficient control URMT cells, neither MNT nor MAX were bound to the *MNT* promoter. In the same way, MNT and MAX were bound to the positive control *Atrogin-1* promoter in URMax34 but not in URMT control cells (Figure 4B). These data show that MNT most likely binds to its own promoter as a heterodimer with MAX and suggest a possible negative regulation of MNT own expression.

### MNT localizes in the cytoplasm of MAX-deficient cells

We next examined the subcellular localization of the excess MNT present in MAX-deficient cells. URMT and URMax34 cells were treated with Zn^2+^ to induce MAX and we performed a cytoplasmic and nuclear fractionation to analyse MNT levels. In control URMT cells, MNT protein was found in the nuclear fraction, as expected, but also at similar levels in the cytoplasm. In contrast, in URMax34 expressing MAX, MNT protein was found only in the nucleus (Figure 5A), the localization typically found for MNT (*Lafita-Navarro et al 2016*). We were not able to use untreated URMax34 cells as a control as they express some MAX even in the absence of Zn^2+^ (Figure 1B). Cytoplasmic MNT was also observed in H1417 cells, human cells deficient in MAX. As a control, the localization of MNT was analysed in HEK293T cells, that express the MAX protein. Cytoplasm/nucleus fractionation revealed that MNT and MAX were localised in the nucleus but not in the cytoplasm of HEK293T cells (Figure 5A). To test if the localization of MNT depended on MAX, we silenced the MAX protein in K562 cells with siRNA and we carried out cytoplasm/nucleus fractionation. In control K562 cells, MNT and MAX were mainly localised in the nucleus. In contrast, in MAX-depleted cells, a significant amount of MNT was found in the cytoplasm of K562 cells (Figure 5B). It is noteworthy that nuclear MNT levels were similar in URMT and URMax34, as well as in control K562 and MAX-depleted K562 cells. As controls for nuclear proteins, we used MYC and CTCF (Figure 5B). Taken together, the data indicates that MAX absence leads to an upregulation of MNT protein and that the excess of MNT accumulates in the cytoplasm.

**Figure 5.**
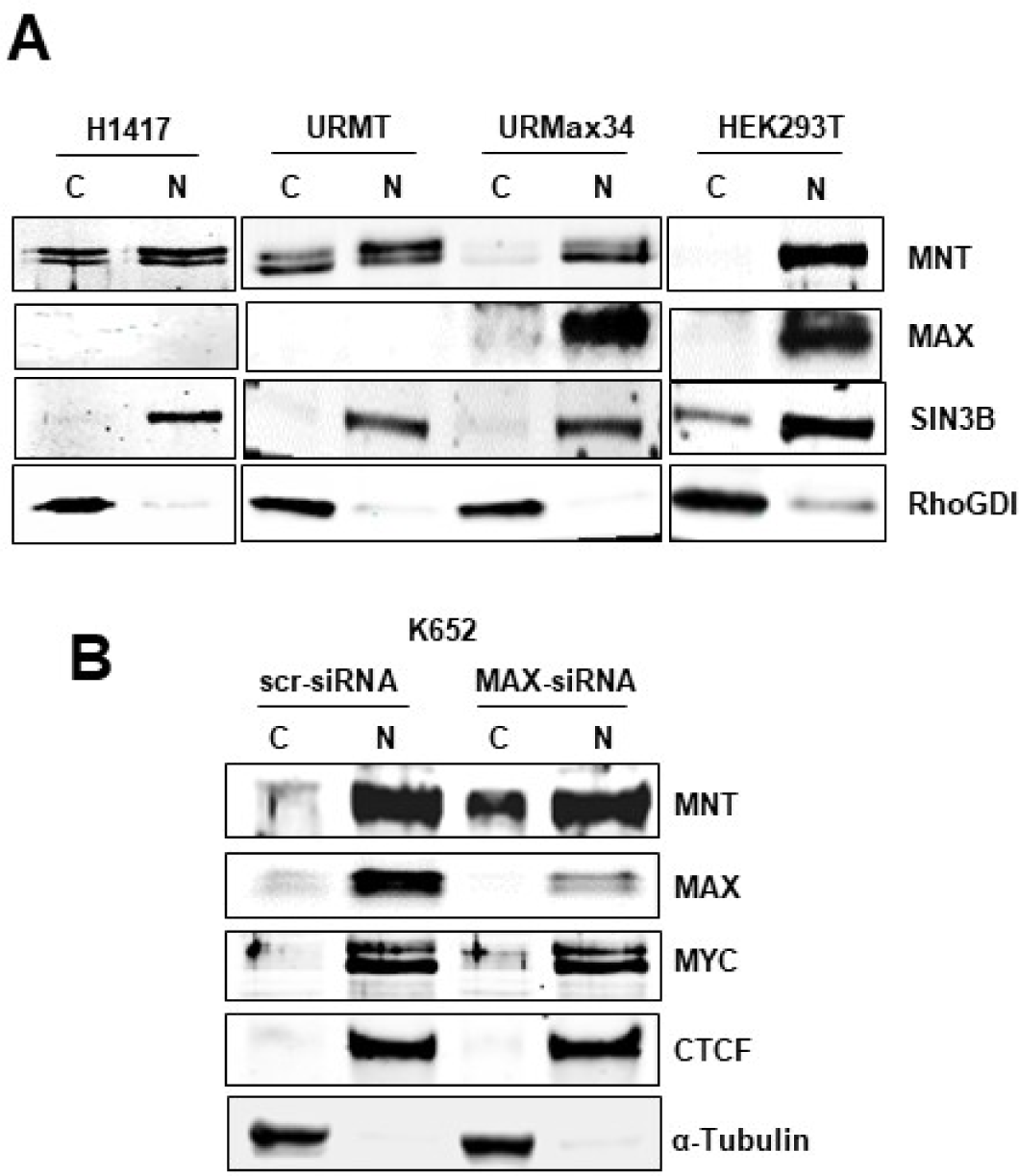
MNT subcellular localization depends on MAX. (A) Cell extracts were subjected to cytoplasmic/nuclear fractionation in H1417 (human lung cells deficient in MAX), HEK293T, URMT and URMax34 treated for 24 h with 1 00 μM~Zn^2+^ to induce MAX. The levels of MNT and MAX were determined in each fraction by immunoblot. SIN38 and RhoGDI (ARHGDIA) were used as nuclear and cytoplasmic markers respectively. “C” refers to cytoplasmic fraction and “N” to nuclear fraction. Similar results were obtained in three biological replicates. (B) K562 were transfected with siRNA against the *MAX* gene and 48 h later cell extracts were prepared and subjected to cytoplasmic/nuclear fractionation. The expression of MAX, MNT and MYC were analyzed by immunoblot. The expression of CTCF and α-tubulin were analyzed as a control for nuclear and cytoplasmic proteins respectively. scrRNA, control scrambled siRNA. The results were reproduced in three biological replicates.

### MNT knock-down impairs cell proliferation in MAX-deficient cells

We next asked for a possible biological effect of MNT on cell proliferation in the UR61 MAX-deficient cells. We decided to knock-down MNT through siRNAs. For this, we used lentiviral particles containing two short-hairpin constructs against the rat MNT gene. Two shMNT constructs that efficiently reduced MNT levels were used, termed shMNT-1 and shMNT-2 (Figure 6A). The vectors also carried a puromycin-resistance gene. UR61 cells were transiently co-transfected with the shMNT constructs (or the empty vector) and a GFP expression vector in a proportion 1:5 to ensure that the GFP-positive cells also incorporated the shMNT plasmid. Six days after transfection, the fraction of GFP positive cells were analysed by flow cytometry. The results showed that the fraction of GFP positive cells was clearly reduced in cells with depleted MNT (Figure 6B), suggesting that MNT loss resulted in impaired cell proliferation. In a second approach, we transfected the UR61 cells with the sh-MNT constructs and counted viable cells after 3 and 7 days of transfection. As shown in Figure 6C, MNT-silenced UR61 cells grew slower than controls. We also performed clonogenic assays in UR61 transfected with the shMNT and/or a MAX expression vector (in 1:3 proportions to ensure that the shMNT containing cells had also incorporated the MAX vector) as well as the empty vectors. 24 h after transfection puromycin was added and after selection the colonies were stained with crystal violet, the dye was solubilised and quantified. The results showed that MNT depletion provoked a dramatic growth inhibition. These results of MNT depletion impairing cellular proliferation in a MAX-independent manner are unprecedented. These results also showed that UR61 cells overexpressing the MAX protein grew more slowly than the control (Figure 6D), confirming the effect previously reported for the MAX-deficient PC12 pheochromocytoma cells (UR61 parental cells) (*Hopewell & Ziff 1995*). In addition, when MNT silencing was accompanied by MAX enforced expression, the inhibition of UR61 cell proliferation was stronger (Figure 6D). Consistent with the anti-proliferative effects of MNT depletion, we failed to generate stable MNT-silenced UR61 cell lines (not shown). In order to generate UR61 cells with MNT overexpression we transfected UR61 with MNT expression vectors but we were not able to detect a significant increase in MNT protein levels upon transfection. This is likely due to the autoregulation of MNT protein levels, as the overexpression of MNT protein was readily detected by immunoblot when the transfected cells were treated with the proteasome inhibitor bortezomib (Figure 6*-figure* s*upplement 1A*). In contrast, some increase in the levels of MNT was detected when MNT was transfected in MAX-expressing URMax34 cells (Figure 6*-figure* s*upplement 1B*) or when MNT was co-transfected with MAX in UR61 cells (Figure 6*-figure* s*upplement 1C*).

**Figure 6.**
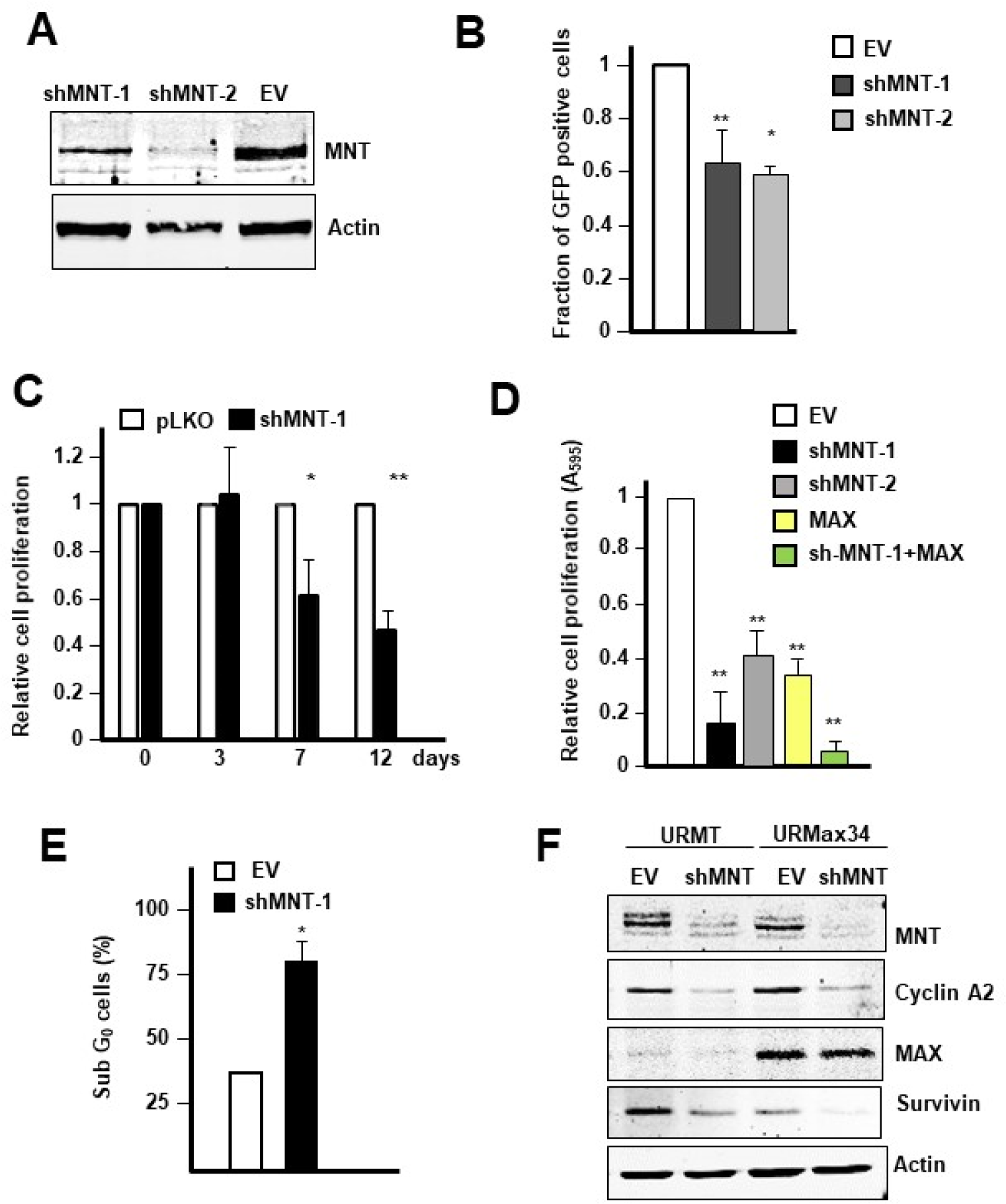
M NT kn oc k-d own impairs cell pro liferation in MAX-deficient cells. (A) Silencin g of M NT by short-hairpin constructs. URMT cells were transfected with vectors encoding sh-MNT-1 and shMNT-2. 24 h after transfection the cells were treated with puromycin (0,3 μg/ml) and 72 h after transfection the cells were lysed and the levels of MNT analyzed by immunoblot. β-actin levels were determined as a control for protein loading. (B) Fraction of GFP-expressing UR61 cells assessed by flow cytometry 7 days after co-transfection with GFP and sh MNT vectors (in proportion 1:5) and analyzed by flow cytometry. The data are shown as mean ± s.e.m. of three independent experiments, \**P* =0.006, \*\**P* = 0.008. (C) Cell proliferation determined by cell counting at 3, 7 and 12 days after transfection of shMNT-1 or the empty vector pLKO1. The data are shown as relative mean values ± s.e.m. of three independent experiments, \**P* = 0.01, \*\**P* = 0.008. (D) Cell growth determined by crystal violet staining in UR61 cells transfected with the indicated vectors. After 15 days of puromycin selection the colonies were stained with crystal violet and the dye was solubilised and quantified by absorbance at 595 nm. EV, empty vector (pLKO for shMNTs and pCEFL for MAX). \*\**P* < 0.005. Data show mean values ± s.d. from three independent experiments. (E) Fraction of sub G_0_ UR61 cells transfected with the shMNT-1 vector and empty vector (EV). Cells were fixed and stained with propidium iodide at day 6 post-transfection and puromycin selection. The percentage of cells containing less than 2C DNA content was determined by flow cytometry. The data are mean values of three independent experiments ± s.d., \**P* = 0.014. (F) Levels of MNT determined by immunoblot in URMT and URMax3 4 cells 72 h after transfection with the shMNT vector and treated for 12 h w ith 100 μM Zn^2+^ MAX, cyclin A2 and survivin (BIRC5) were also determined. β-actin levels were determined for protein loading control. Two experiments were performed with similar results.

Since the depletion of MNT impairs cell proliferation of UR61 cells, we wondered whether this was accompanied by cell death. To study this, the levels of survivin (BIRC5) in URMT and URMax34 were analysed by immunoblot (Figure 6F) and the results showed a decrease in survivin and cyclin A (a marker of proliferation) in cells with depleted MNT, both in MAX-deficient cells (compare lanes 1 vs 2 in Figure 6F) and in MAX-expressing cells (compare lanes 3 vs 4 in Figure 6F), although the effect was stronger in these latter cells. To check the contribution of apoptosis, we also measured DNA content flow cytometry. We found that the fraction of cells with a sub-G0 amount of DNA was higher in cells transfected with the shMNT vector (Figure 6E). The results suggest that the depletion of MNT leads to cell proliferation arrest and apoptosis in UR61.

### MNT modifies gene expression in MAX-deficient cells

Since the data shown above indicated that MNT knock-down in UR61 cells leads to growth arrest and cell death, and considering that MNT is a transcription factor, we set out to study if MNT regulates the transcriptional program in the absence of MAX. For this purpose, we investigated the effects of MNT depletion on the transcriptome of cells with and without MAX. URMT and URMax34 cells were transfected with the short-hairpin RNA construct against the rat MNT gene (shMNT-1) or the empty vector as control in two different biological replicates. We first checked the appropriate conditions to efficiently silence the expression of MNT upon transfection of the shMNT constructs in URMT and URMax34 cells. A significant MNT depletion was consistently achieved at 72 h post-transfection (not shown). Thus, for the RNAseq experiment, the URMT and URMax34 cells were transfected with the shMNT-1 construct and two days after transfection, cells were treated with Zn^2+^ for 24 h to achieve significant MAX-induced expression (in URMax34) as well as significant MNT depletion (Figure 7A). Two independent transfections and RNA preparations of each cell line were submitted for next generation sequencing RNA-seq and the data was processed bioinformatically as escribed in the Methods section.

**Figure 7.**
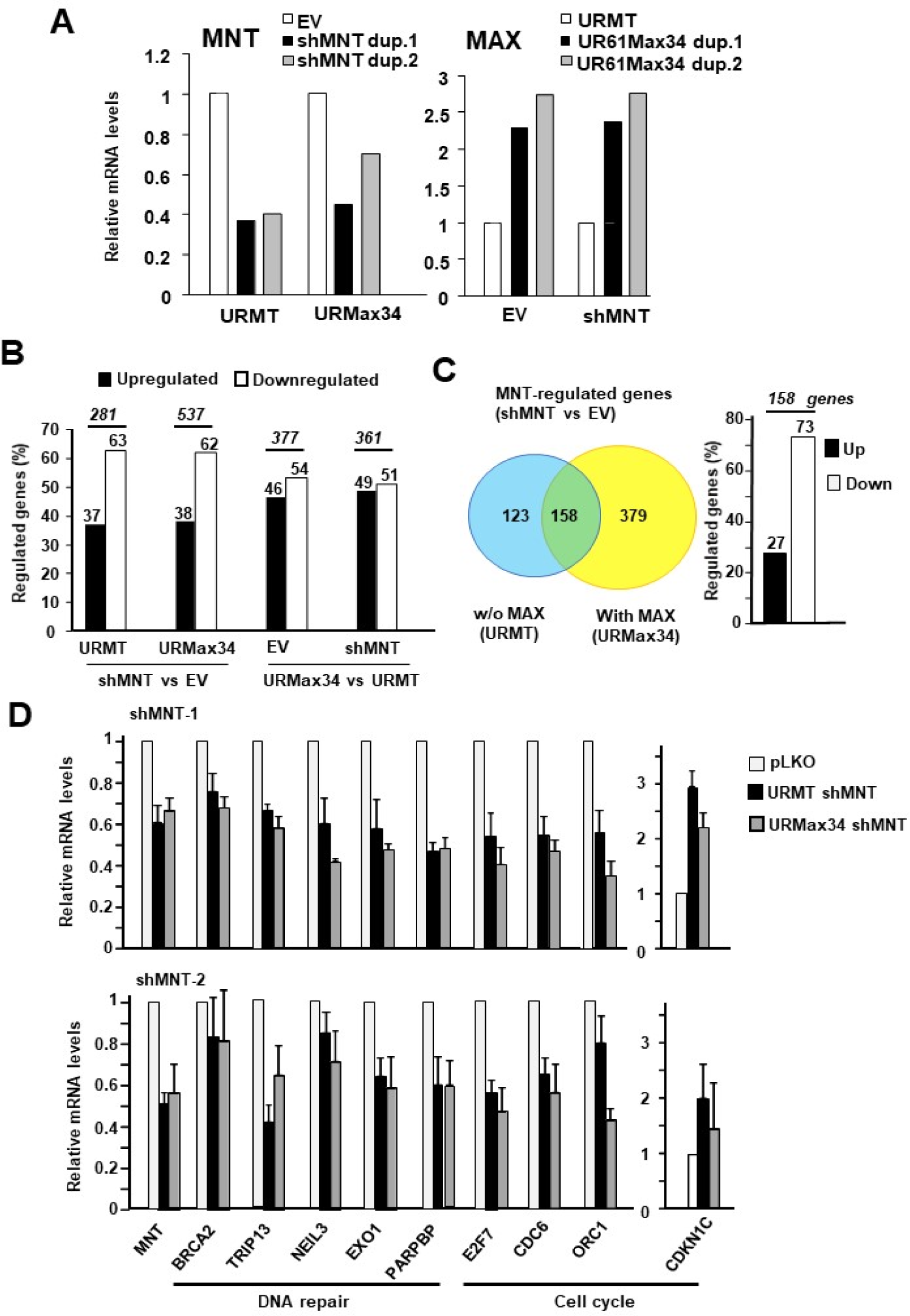
Gene expression changes in MNT knock-down cells. (A) mRNA expression of MNT and MAX in URMT and URMax34 cells lysed 72 h after transfection w ith shMNT and 24 h of treatment with 100 μM Zn^2+^. The experiments were performed in duplicated (dup.1 and dup.2) biological replicates. These RNAs were subjected to RNA-seq. (B) Percentage of upregulated and downregulated genes in cells transfected with th e shMNT or pLKO (empty) vectors in URMT and URMax34 cells, as indicated at the bottom. The number of genes regulated comparing the different samples are indicated at the top of the figure. The gene inclusion criteria were a change in mRNA level ≥ 0.7 or ≤ 0.7 log2 FPKM fold change. T he RNA-seq data are from two independent experiments. The genes regulated in the different comparisons are listed in the *Supplementary Files 1 to 4*. (C) Venn diagram showing the genes that are regulated by MNT in MAX-deficient (URMT) or MAX-expressing cells (URMaX34). The graph at the right shows the number of up- or down-regulated genes in the group of overlapping genes. (D) mRNA expression regulation due to MNT silencing in genes selected after the RNA-seq data analysis. URMT and URMax34 were transfected with shMNT-l (upper panel) and shMNT-2 (lower panel) or empty vector pLKO. After 48 h cells were treated for 24 h with 100 μM Zn^2+^, total RNA was prepared and mRNA levels of the indicated genes were determined by RT-qPCR. Most of the selected genes are involved in DNA repair or cell cycle as indicated at the bottom. The data are represented as mean ±s.e.m. from four (shMNT-1) or three (shMNT-2) independent transfection experiments. In all cases, *P* < 0.05.

Once we obtained the expression values of each experimental replicate we compared the different conditions. For all of the different comparison we grouped the genes that were up-regulated and down-regulated in both biological replicates considering a log2 RPKM fold change higher than 0.7 or smaller than −0.7 of the corresponding control and a p-value < 0.1. The heatmaps of the gene expression signatures clearly showed that the depletion of MNT in both URMT and URMax34 induced gene expression changes (Figure 7*-figure supplement 1*) indicating that MNT can be involved in transcriptional regulation without the concourse of MAX. Specifically, 281 genes were regulated upon MNT depletion in MAX-deficient URMT cells (Figure 7*-figure supplement 2*) and 537 genes in MAX-expressing cells (i.e., URMax34 treated with Zn^2+^) (Figure 7*-figure supplement 3*). Of those, approximately 62 % of the genes were down-regulated upon MNT depletion in both cell lines (Figure 7B). In addition, the URMax34 vs URMT gene expression signatures heatmaps also showed that the expression of MAX induced gene transcriptional changes independently of MNT depletion (Figure 7*-figure supplement 1*). Specifically, 377 genes were found to be differentially expressed when comparing URMax34 and URMT cells (Figure 7*-figure supplement 4*) and 361 genes when comparing URMax34 and URMT cells depleted of MNT (Figure 7*-figure supplement 5*). Among *MAX*-regulated genes, roughly half of the genes were downregulated (Figure 7B). Importantly, the comparison between the lists of differentially expressed genes upon MNT-depletion in cells without MAX (URMT) and with MAX (URMax34) revealed 158 shared genes in the two biological replicates (Figure 7*-figure supplement 6*). This indicated that 56% and 30% of MNT-induced transcriptional changes in each cell line respectively are regulated by MNT in a MAX-independent manner (Figure 7C).

The genes showing expression changes in the MNT-depleted URMT and URMax34 cells were analyzed with the GSEA and the MsigDB platform (http://software.broadinstitute.org/gsea/msigdb). The genes regulated upon MNT knockdown were compared with the gene sets derived from the biological process gene ontology and the comparison showed that genes belonging to cell cycle processes are the most enriched set in MNT-depleted cells both in MAX expressing and not expressing cells (Figure 7*-figure supplement 7*). This functional analysis of the gene transcriptional signatures are consistent with the biological effects of MNT-depletion in cell proliferation.

Thus, in order to explain and confirm the effects of MNT ablation on UR61 cell growth, we selected several genes involved in cell cycle and DNA replication which expression was changed according to the RNA-seq analysis. URMT and URMax34 cells were transfected with the two shMNT constructs and the empty vector following the same conditions as the conditions used for the RNAseq experiment. The expression of the selected genes was analyzed by RT-qPCR. Figure 7D shows that genes involved in cell cycle and DNA replication (*E2F7*, *CDKN1C*, *CDC6*, *ORC1*) and, DNA repair (*BRCA2*, *TRIP13*, *NEIL3*, *EXO1*, *PARPB*), were down-regulated upon silencing of *MNT* with the two shMNT constructs, except *CDKN1C* (p57, a cell cycle inhibitor) which was up-regulated. Altogether, the results are consistent with the negative effect of MNT depletion on cell proliferation in UR61 cells (Figure 6).

We then wondered how could MNT act as a transcription factor in cells lacking MAX. Besides MAX, MNT can also bind the HLH protein MLX (*Cairo et al 2001, Meroni et al 2000*). To show if this interaction also takes place in UR61, we transfected MLX-Flag into URMT cells and 48 h later lysates were immunoprecipitated with anti-MLX antibody and anti-MNT antibodies. The immunoprecipitates were analysed by immunoblot and the results showed the interaction between MNT and MLX (Figure 8A). We next studied this interaction in URMax34 cells treated with Zn^2+^, i.e. cells expressing MAX. The immunoblot results showed that MNT and MLX also interacted but the interaction was weaker when MAX expression was induced by Zn^2+^. Thus, the data suggest that, at least in our experimental conditions, MNT-MAX dimers were formed more efficiently than MNT-MLX dimers (Figure 8B). To confirm the MNT-MLX interaction in URMT cells, we prepared a HA-tagged MNT mutant with a deletion of most of the HLH domain of mouse MNT, termed ΔHLHMNT-HA (Figure 8C). This mutant or the wild-type counterpart was transfected into URMT and the corresponding lysates were immunoprecipitated with anti-HA. The results showed that HLH domain is required for the MNT-MLX interaction (Figure 8D).

**Figure 8.**
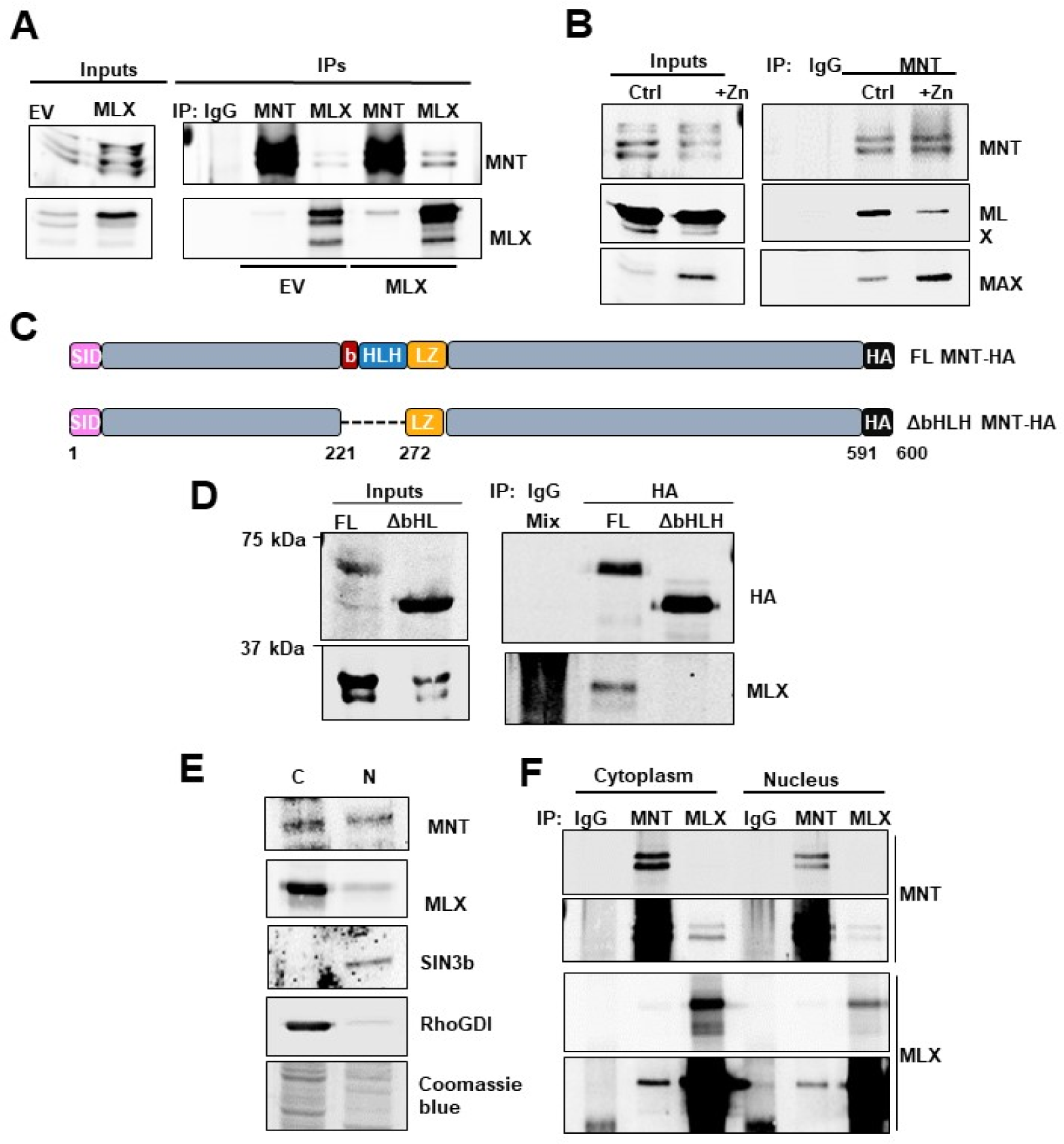
MNT and MLX interaction in UR6 1 cells. (A) URMT cells were transfected with a MLX expression vector or the empty vector (EV). 48 h after transfection, total cell Iysat es were prepared and precipitated with anti-MNT or anti-MlX antibodies, as well as unspecific IgG. The presence of MNT and MLX were detected in the immunoprecipitates by immunoblot T he results w ere reproduced in two experiments. (B) UR Max34 cells were transfected with a MLXexpression vector an d 48 h after transfection cells were treated with 100 μM Zn^2+^ for 24 h. Then total cell Iysates were prepared and immunoprecipitated with anti-MNT. As a control, a mixture of Iysates from cells treated and untreated with Zn^2+^ was immunoprecipitated with unspecific IgG. The presence of MNT, MLX and MAX in the immunoprecipitates were analyse d by im munoblot. The results were reproduced in three experiments immunoprecipitations. (C) Schematic representation of the full-length (FL) and the deletion mutant ΔHLH MNT-HA used in subsequent experiments. The SID (Sin3 Interacting Domain), bHLH, LZ domains, HA tag and amino acids of the murine protein are indicated. (D) URMT cells were co-transfected with a vector expressing MLX and the constructs shown in (C) as indicated at the top. 48 h after transfection, cell Iysat es were prepared and the cells were immunoprecipitated with mouse anti-HA antibody to pull down the exogenous MNT proteins constructs. As a control, Iysates (a mixture of Iysates from cell transfected with FL MNT and ΔHLH MNT) were also immunoprecipitated with unspecific IgGs. The immunoblot analysis using a rat monoclonal anti-HA antibody revealed the presence of FL MNT and the smaller “ΔHLH MNT The presence of MLX in the immunoprecipitates was analysed with an anti-MLX antibo dy. (E) MLX localization in UR61 cells. Nuclear (“N”) and cytoplasmic (“C”) extracts were prepared from URMT cells as described in Materials and Methods and the levels of MNT and MLX were determined in each fraction by immunobloL SIN3B and RhoGDI (ARHGDIA) were used as nuclear and cytoplasmic markers respectively. The antibodies used are as described in the *Supplementary File 1*. A picture of the gel stained with Coomasie Blue after transference is also shown as an indicator of the total amount of proteins in the cytoplasmic and nuclear fraction. (F) Interaction between MNT and MLX in the nucleus and cytoplasm. URMT cells were transfected with a MLX expression vector and 48 h later nuclear and cytoplasmic extracts were prepared and immunoprecipitated with anti-MNT or anti-MlX antibodies. The levels of MNT and MlX in th e immunoprecipitates were assayed by immunoblot. The results were reproduce in two independent experiments.

In contrast to MAX, MLX is expressed in both nucleus and cytoplasm (*Billin et al 2000*). By immunoblot analysis it was found that MLX was present in nucleus and, at a lesser extent, in the cytoplasm of URMT cells (Figure 8E). Next, we prepared nuclear and cytoplasmic fractions of URMT cells and studied by immunoprecipitation with anti-MNT and anti-MLX antibodies if MNT and MLX interact. The results showed that this was indeed the case in both nuclear and cytoplasmic fractions (Figure 8F). The proximity ligation assay also showed that MNT and MLX interacted in both nucleus and cytoplasm (not shown).

Finally, it is described that MNT can form homodimers in two-hybrid experiments (*Hurlin et al 1997, Meroni et al 1997*). Thus, a possibility is that MNT acts a transcription factor as a homodimer. This homodimerization has not been demonstrated *in vivo* so we tested it in HEK293T and UR61 cells. We first co-transfected HEK293T cells with GFPMNT and Flag-MNT constructs and immunoprecipitated with anti-GFP antibody. The immunoblot analysis demonstrated the presence of the smaller Flag-MNT protein in the material immunoprecipitated with anti-GFP (Figure 9A). As expected, both big and small MNT forms were detected when the immunoblots were analysed with anti-MNT antibody (Figure 9A). The results suggested the presence of homodimers between GFP-MNT and MNT-Flag in HEK293T cells. As a control, the lysates with anti-MAX antibody were imunoprecipitated, and both MNT and GFP-MNT were found to be bound to MAX (not shown). Next, we investigated the MNT homodimerization in the UR61 system. URMT cells were infected with lentiviral particles containing the GFP-MNT construct, immunoprecipitated, and the lysates analysed by immunoblot with anti-GFP antibody and anti-MNT antibodies. The results showed that endogenous MNT was present in the immunoprecipitates with anti-GFP (Figure 9B). The results suggest that MNT forms homodimers in human HEK293T cells and in MAX-deficient rat URMT cells. As MNT dimerization in yeast two hybrid assays depends on the HLH-LZ (*Hurlin et al 1997, Meroni et al 1997*), we asked whether HLH was involved in the homodimerization *in vivo*. We transfected the MNT mutant lacking the bHLH region (ΔbHLH MNT) (Figure 8C) as well as the wild-type form and we carried out immunoprecipitations with anti-HA antibodies which should only recognize the exogenous proteins. The immunoblot analysis of the immunoprecipitates revealed that ΔbHLH MNT was unable to bind GFPMNT (Figure 9C). The same experiment was performed in URMT cells and the same result was observed, i.e, GFP-MNT bound to wild-type MNT-HA but not to the ΔbHLH form (Figure 9D). We conclude MNT homodimerize through the HLH domain, as expected. Altogether, the results indicate that in URMT cells, MNT can form homodimers or heterodimers with MLX, whereas in URMax34 (in the presence of Zn^2+^), MNT can also form heterodimers with MAX, as schematized in Figure 9E.

**Figure 9.**
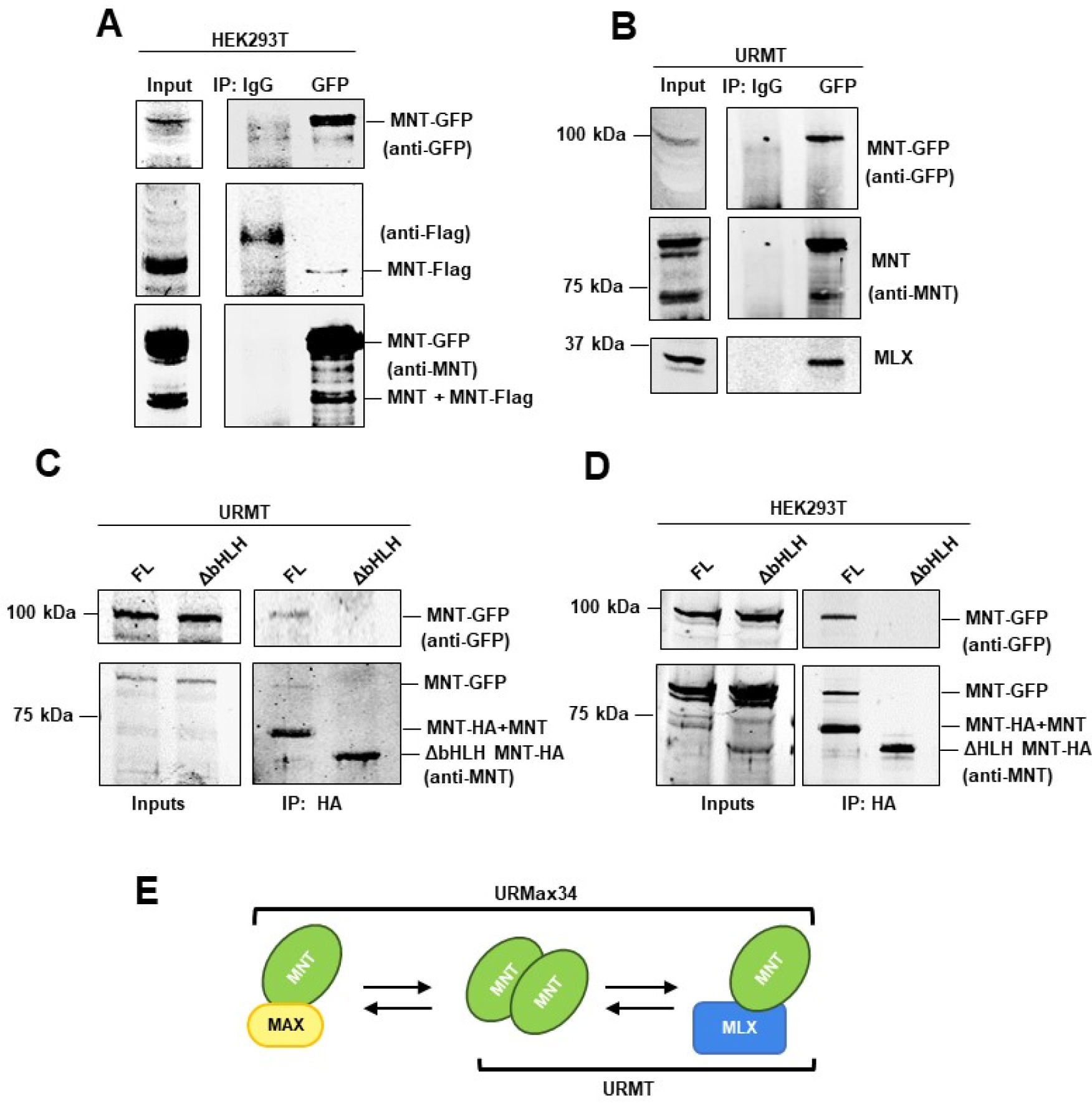
MNT homodimerization in UR61 cells. (A) HEK293T cells were co-transfected with a GFP-MNT and MNT-Flag expression vectors, and 24 h later Iysates were prepared and immunoprecipitated with anti-GFP antibody. The presence of MNT-GFP and MNT-Flag were revealed by immunoblotting the antibodies indicated at the right. As a control, Iysates were precipitated with unspecific IgG. The antibodies used in each immunoblot are indicated at the right The results were reproduced in two immunoprecipitations. (B) URMT cells were infected with lentivirus encoding a MNT-GFP an d 72 h later were transfected with MNT-Flag and treated with 15 nM bortezomib for 12 h before harvesting. 48 h after transfection Iysates were prepared and immunoprecipitated w ith anti-GFP antibody. The presence of MNT-GFP, MNT and MLX in the immunoprecipitates were assessed by immunoblot The antibodies used in each immunoblot are indicated at the right. The results were reproduced in two experiments. (C) URMT cells were infected with lentivirus encoding a GFP-MNT and 72 h later transfected with expression vectors for full length (FL) MNT or ΔHLH MNT (Figure 8C) as indicated at the top of each lane. 48 h after transfection, cell Iysates were prepared and immunoprecipitated with anti-HA to pull down the transfected MNT proteins. The levels of MNT-GFP and MNT were determined by immunoblot using anti-GFP and polyclonal anti-MNT antibody. (D) HEK293T cells were co-transfected with expression vectors for MNT-GFP and expression vectors for FL MNT or ΔHLH MNT (Figure 8C) as indicated at the top of each lane. 24 h later Iysates were prepared and immunoprecipitated with anti-MNT antibody. The levels of MNT-GFP and MNT were determined by immunoblot using anti-GFP and polyc lonal anti-MNT antibody. The results were reproduced in two experiments. (E) Schematic summary of the interactions of MNT found in URMT (MAX-less) and URMax34 (expressing MAX when treated with Zn^2+^).

## DISCUSSION

In this study, we report several novel findings: (i) MNT is required for optimum proliferation even in the absence of MAX; (ii) in the absence of MAX, MNT expression is elevated and a significant fraction localizes in the cytoplasm; (iii) MNT represses its own transcription in a MAX-dependent manner; (iv) MNT is able to regulate the expression of genes in the absence of MAX and (v) MNT forms homodimers in vivo. Previous reports suggest that MNT functions as a “MYC buffer”, curbing excessive MYC activity. For instance, *in vivo* MNT knockdown impedes MYC-driven lymphomagenesis (*Campbell et al 2017*) whereas MNT silencing leads to MYC-like phenotypes (*Hurlin et al 2003, Nilsson et al 2004, Walker et al 2005*). Our data are in line with this critical MNT function, as they show a tight control of MNT expression by which MNT limits its own mRNA expression. MNT protein levels are also under strict control in UR61 cells, as MNT cannot be overexpressed at high levels upon transfection unless the proteasome is inhibited. Given the relevance of MNT to modulate MYC activity and its central position between the MYC-MAX and MLX-MONDO networks, the activities and regulation of MNT are a key issue on MYC-dependent oncogenesis.

MNT expression was high in some MAX-deficient cells like the rat UR61 cells and human lung carcinoma H1417 cells. The absence of MAX was in part responsible for this effect because (i) MAX re-expression in UR61cells results in decreased MNT mRNA and protein, and (ii) MAX overexpression in K562 cells results in MNT down-regulation whereas MAX knock-down results in MNT up-regulation. The mechanism for this MAX effect on MNT levels depended, at least partially, on the auto-repression of MNT expression by MNT-MAX dimers. Luciferase reporter experiments showed that MNT represses its own promoter (which contains two conserved E-boxes) and ChIP assays showed that MNT binds to its own promoter when MAX is ectopically expressed in UR61 cells. In agreement with these results, significant levels of MNT localize to the cytoplasm of MAX-deficient cells whereas MNT remains nuclear in MAX-expressing cells. This would explain why MAX re-expression leads to a decrease in MNT levels: MNT-MAX dimers would be formed to repress MNT expression in the nucleus. On the contrary, in the absence of MAX, MNT is expressed at higher levels since there would be no negative MNT autoregulation. The model is schematized in Figure 10.

**Figure 10.**
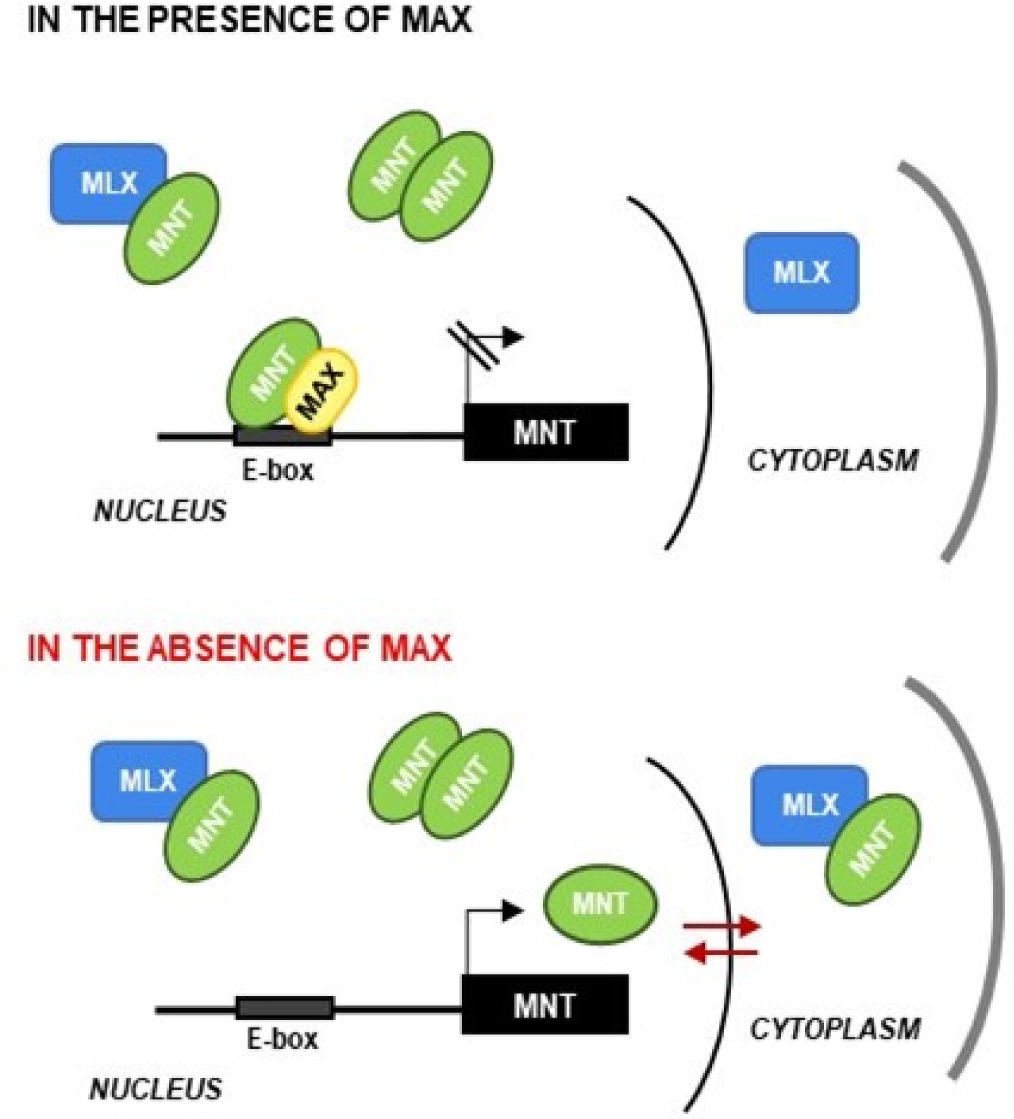
Model of MNT fates depending on MAX. In MAX-expressing cells, most of the MNT is retain ed in the cell nucleus where it limits its own expression. In MAX-deficient cells as UR61, MNT is distributed in nucleus and cytoplasm and is unable to bind the promoter and regulate its transcriptional activity. The model includes the presence of MNT-MNT homodimers and the interaction MNT-MLX in the cytoplasm, which might be responsible for the MNT partial localization in the cytoplasm in MAX-deficient cells.

We explored possible MAX-independent effects of MNT using the UR61 model. Although originally MAX was defined as an obligate dimerization partner of MYC, work carried out in the PC12 model and in *Drosophila* indicated that MYC can function in a MAX-independent manner, for example in inhibition of differentiation (*Gallant 2013, Maruyama et al 1987, Vaque et al 2008*). MYC overexpression blocks RAS-induced differentiation of UR61 cells (*Vaque et al 2008*) however, we did not detect a significant effect of MNT depletion on differentiation (not shown). In contrast, depletion of MNT in MAX-deficient UR61 cells impairs cell proliferation. This is the first report on a MAX-independent function of a MXD protein. In contrast, in rat and mouse fibroblasts, MNT knock-out leads to increased proliferation and transformation capacities (*Hurlin et al 2003, Nilsson et al 2004*). The fact that MNT depletion impairs UR61 proliferation (in a MAX-independent manner) adds complexity to the MNT-MYC functional interactions. Actually, in gastric cancer high MNT expression correlates with shorter survival whereas MYC overexpression has the opposite effect (http://kmplot.com/analysis/).

Since MNT is a transcription factor, we compared the transcriptomes of parental UR61 cells versus cells with depleted MNT. Upon MNT depletion, a number of genes involved in cell cycle progression and DNA damage response were downregulated in cells with and without MAX. This is concordant with the decrease in proliferation exerted by MNT silencing in UR61. It is open to discussion how MNT regulates genes in the absence of MAX. MNT can form heterodimers with the HLH protein MLX (*Cairo et al 2001, Meroni et al 2000*) and we have confirmed the ability of MNT to bind MLX in UR61 cells, but when MAX is expressed, the dimers MNT-MAX are favoured versus MNT-MLX. As a relevant fraction of MLX is cytoplasmic and we have shown that MNT and MLX also interact in the cytoplasm, this interaction could help to explain the increase in cytoplasmic MNT observed in MAX-depleted cells. The model is depicted in the Figure 10. Moreover, we have shown that MNT homodimerizes in UR61 cells, and therefore it is likely that MNT regulates genes as a homodimer. Further work is required to clarify this point. The results reported here show a strict autoregulation of MNT, supporting a pivotal role of MNT in the control of cell proliferation even in the absence of MAX. Moreover, we showed that MNT has activities in the absence of MAX. Given that MNT participates in both the MAX-MYC and MLX-MONDO networks, the regulation of MNT is a relevant issue in cell biology and in tumor development promoted by MYC.

## MATERIALS AND METHODS

### Cell lines and transfections

Cell lines were obtained from ATCC and grown in either RPMI-1640 or DMEM (Lonza) supplemented with 10% fetal bovine serum (Lonza), 150 µg/ml of gentamicin and 2 µg/ml of ciprofloxacin. To generate the URMT and URMax34 cell line, UR61 cells (*Guerrero et al 1988*) were electroporated (260 V, 1 mFa, BioRad apparatus) with a pHeBo-MT or pHeBo-MT-Max vector which carries a human MAX cDNA under the control of metallothionein promoter (*Canelles et al 1997*). Cells were selected with 0.2 mg/ml of hygromycin (Life Technologies) and cell clones were isolated by limiting dilution Transient transfections were carried out using the Ingenio Electroporation solution (Mirus) in an Amaxa nucleofector. The cells were transfected or transduced with expression vectors for MAX (pCEFL-MAX(*Mauleon et al 2004*), MLX (β isoform, (pMS18-MLX,(*Billin et al 1999*), human MNT (pCMVSport6-MNT, Origene Technologies), MNT-Flag (human MNT with FLAG at the C-ter lentiviral Lv158 vector Genecopoeia), MNTGFP (human MNT with GFP at the N-ter, lentiviral Lv103 vector Genecopiea,), FL MNT-HA (full-length murine MNT tagged at the C-ter with hemagglutinin epitope, HA), ΔHLH MNT-HA (murine MNT carrying a deletion of amino acids 221-272 amino acids and tagged with HA) (both in pcDNA3 with the zeocin resistance gene inserted), short-hairpin human MNT-1 (shMNT-1, pLKO-shMNT from Sigma Mission, TRCN0000085733), or MNT-2 (shMNT-2, pLKO-shMNT from Sigma Mission, TRCN0000235815), siRNA for human MAX (Sigma, SASI_Hs01_00011941).

### Immunoblot and immunoprecipitation

Cell lysis, immunoblots and immunoprecipitations were performed as described.(*Garcia-Sanz et al 2014*). The antibodies used are described in *Supplementary File 1*.

### Luciferase reporters and assays

To generate the 850MNT-luc, 220MNT-luc, 570MNT-luc reporter vectors, two pair of primers were designed for each construct (*Supplementary File 2*), targeting sequences of the human genome corresponding to 850 bp upstream the transcription start site (TSS) of *MNT* gene. The amplified DNA was inserted into the pBV-luc reporter vector at the EcoRV and HindIII sites (*He et al 1998*). The sequence of MNT promoter was from the UCSC genome browser (http://genome.ucsc.edu/; GRCh37/hg19). Cells were transfected with jetPEI reagent (Polyplus) and luciferase assays were performed as described (*Garcia-Sanz et al 2014*).

### Cell proliferation and cell cycle analysis

Cell proliferation was monitored with a cell counter (NucleoCounter NC-100, Chemometec) or a cytometer (Guava PCA, Merck Millipore). For the clonogenic assays, 1-2 x 10^6^ cells/ml were seeded in a 6-well or 60 mm plate after transfection by electroporation. 48 h post-transfection, cells were selected with puromycin at 0.2-0.5 µg/ml final concentration. After 8-17 days the cells were stained with crystal violet and the dye measured by absorbance at 595 nm as described (*Ferrandiz et al 2009*). To determine the sugG0-G1 population, cells were transfected and 7 days post-transfection the DNA concentration was analyzed by propidium iodide staining and flow cytometry as described (*Albajar et al 2011*).

### RNA analysis

For qPCR, total RNA was isolated using the TRI Reagent^®^ Solution (Invitrogen). cDNA was generated by reverse transcription (RT) using the iScript (Bio-Rad). Quantitative polymerase chain reaction (qPCR) was performed with specific primers (*Supplementary File 2*) using the iTaq™ Universal SYBR^®^ Green Supermix (Bio-Rad) and CFX ConnectTM Real-Time PCR Detection System (Bio-Rad). RNA was converted into cDNA and analysed as described (*Bretones et al 2011*). Levels of mRNA were normalized against actin and RPS14 mRNA levels. For RNA-seq, total RNA was isolated with the RNeasy kit (Qiagen) from two independent experiments of MNT silencing in URMT and URMax34 cells, both treated with 100 µM ZnSO_4_ for 24 h. mRNA libraries were prepared using the Illumina TruSeq RNA Sample Prep Kit v2 (kit RS-122-2002, Illumina). A minimum of 40 million 50 bases single-end reads per sample were obtained. Tophat algorithm (*Trapnell et al 2009*) was used to align the data using a set of gene model annotations and/or known transcripts of rat genome (rn4) obtained from RefSeq database. Cufflinks software (*Trapnell et al 2010*) was run to estimate transcripts abundances represented in RPKM units (reads per kilobase per million reads) as described (Mortazavi et al 2008). The gene expression of the genes are represented as RPKM values (Figure 8 *–Supplement 2 to 5*).

### Chromatin immunoprecipitation

Fractionation of total cell extracts into cytoplasmic and nuclear fractions and chromatin immunoprecipitation (ChIP) were performed essentially as described (*Garcia-Sanz et al 2014*). Immunoprecipitated DNA was purified with the QIAquick PCR Purification Kit (Qiagen) and analysed by qPCR. The antibodies and primers used are described in *Supplementary Files 1 and 2,* respectively.

### Nuclear/cytoplasmic fractionation

Cytoplasmic and nuclear extracts were prepared essentially as described (*Conacci-Sorrell et al 2010*). The hypotonic buffer to lyse cells was 10 mM HEPES pH 7, 10 mM KCl, 0.25 mM EDTA pH 8, 0.125 mM EGTA pH 8, 0.5 mM spermidin, 0.1% NP40, 1 mM DTT and proteases inhibitors The hypertonic buffer for nuclear extracts was 20 mM HEPES, 400 mM NaCl, 0.25 mM EDTA, 1.5 mM MgCl_2_, 0.5 mM DTT and proteases inhibitors.

### Statistical analysis

Student’s t-test was used to evaluate the significance of differences between control and experimental groups. A p-value of less than 0.05 was considered as significant. The threshold for expression changes in the RNA-seq analysis was set as log2(RPKM fold change) ≥ 0.7 or ≤ −0.7 and a p-value <0.1.

### Bioinformatic analysis

Genomic sequences of 1000 bp upstream TSS of human, mouse and rat MNT genes were obtained from *H. sapiens* (GRCh37/hg19), *M. musculus* (GRCm38/mm10) and *R. norvegicus* (RGSC 5.0/rn5) assemblies. For the ChIP-seq ENCODE analysis, the data for MAX, MYC and MXI1 was obtained for the indicated cell lines. The SYDH ENCODE project was chosen for representation of the binding peaks in the promoter of the human MNT gene. The sequences corresponding to those peaks were analysed confirming the presence of the E-boxes identified in the human MNT promoter

**Source data**: the RNA-seq data is loaded in the European Nucleotide Archive with the accession number PRJEB23604

## ACKNOWLEDGEMENTS

We are grateful to Montserrat Sánchez-Céspedes and Don Ayer for cell lines and constructs, Maralice Conacci-Sorrel for comments on the manuscript, and Sandra Zunzunegui and Maria Aramburu for technical help.

## AUTHOR ORCID

Javier León, https://orcid.org/0000-0001-5803-0112

